# Antisense ncRNAs during early vertebrate development are divided in groups with distinct features

**DOI:** 10.1101/2020.02.08.940148

**Authors:** Sanjana Pillay, Hazuki Takahashi, Piero Carninci, Aditi Kanhere

## Abstract

Long non-coding RNAs or lncRNAs are a broad class of non-protein coding RNAs that are >200nucleotides in length. A number of lncRNAs are shown to play an important role in gene expression regulation. LncRNAs antisense to a protein-coding gene can act either as positive or negative regulators of overlapping protein-coding mRNAs. Almost 50% of lncRNAs present during development of vertebrates such as zebrafish are of antisense lncRNA class. However, their role in gene expression regulation during development remains enigmatic. To understand the role of antisense lncRNAs in early vertebrate development, we took a computational biology approach to analyze existing as well as novel dataset. Our analysis of RNA sequencing data from zebrafish development indicates that antisense RNAs can be divided into two major classes based on their positive or negative co-expression patterns to the sense protein-coding genes. The ones with negative co-expression patterns or group-1 are maternal antisense lncRNAs that overlap mainly developmental genes. Group-2 with positive expression pattern overlap mainly house-keeping genes. Group-1 antisense lncRNAs are longer and more stable as compared to antisense lncRNAs in group-2. In addition, to answer if antisense RNAs in the two groups are differently localized in cell compartments, we deep-sequenced RNA from cytoplasmic and nuclear compartments during early developmental stages. The analysis of these compartment specific datasets revealed group-1 lncRNAs are cytosolic. Based on the cytosolic nature of group-1 RNAs and their higher complementarity to the overlapping developmental mRNAs, we speculate that the group-1 RNAs might function similar to microRNAs in silencing spurious expression of developmental genes. Group-1 and group-2 RNAs are also distinct in terms of their genomic configuration, conservation, length and transcriptional regulation. These results are not only important in understanding the role of antisense RNAs in development but also for predicting the nature of association between antisense lncRNA and overlapping protein-coding genes.

## INTRODUCTION

Through deep sequencing several novel transcripts are detected which do not code for proteins. Some of these non-protein-coding RNAs or ncRNAs perform housekeeping functions and are expressed constitutively. These include ribosomal RNA (rRNAs), transfer RNAs (tRNAs), small nuclear RNA (snRNAs). However, several other ncRNAs are often expressed in time and space dependent manner. A number of examples suggest that they are involved in regulating gene expression in cells. These cell specific regulatory RNAs are normally divided in to small (< 200b) and long ncRNAs (> 200b). The functional mechanisms of small ncRNAs, such as microRNAs and piRNAs, are well studied (1). Long non-coding RNAs (lncRNAs), on the other hand, still remain enigmatic. LncRNAs can be divided into two broad classes long intergenic RNAs (lincRNAs), which are expressed from intergenic regions between protein-coding genes and non-coding transcripts that overlap a protein-coding gene in antisense direction (AS). In recent years, a lot of efforts have been concentrated on understanding long intergenic RNAs. However, functional mechanisms for a majority (96%) of lncRNAs often remains elusive (2). The FANTOM3 consortium in 2005 reported that up to 72% of potential transcriptional units are transcribed in both directions in mice (3). Studies in humans suggest that AS, such as *Kcnq1ot1* (4), *Airn* (5) and *HOTTIP (6)* are involved in important cellular processes such as imprinting and spatiotemporal coordination of expression during vertebrate development (6). However, for majority of AS, functional mechanism and relationship to the overlapping protein-coding transcript during development remains largely unclear. Previous transcriptomics studies have identified large number of lncRNAs during embryonic development in vertebrates such as zebrafish (7-9). Pauli et al., e.g., identified 1133 lncRNAs across eight stages of zebrafish development, out of these 397 were intergenic lncRNAs, 184 intronic overlapping lncRNAs and the rest 566 were classified as exonic overlapping antisense lncRNAs (8). The abundance of AS lncRNAs during early development combined with functional studies on selected AS supports their importance in early embryonic development (8,10,11).

Zebrafish is one of the popular animal models which is routinely used to understand early vertebrate development (12-14). In Zebrafish, embryonic development starts by fertilization of externally laid eggs and spans across a period of 3 days post-fertilization (dpf). Initially, the embryo undergoes 10 rapid and asynchronous cell divisions which is followed by lengthening of the cell cycle. During development the vertebrate embryo is in a transcriptionally inactive state for a few initial cell divisions. As result, during this period of inactive genome early development of the embryo is completely dependent on maternally provided products. As development progresses, the transcription of zygotic genome is triggered and simultaneous clearance of maternal RNAs and proteins leads to their replacement with newly synthesized zygotic RNAs. This process is called maternal-to-zygotic transition (MZT) or zygotic genome activation (ZGA). In zebrafish, MZT coincides with mid-blastula transition (MBT) and occurs at cell cycle 10 or 3 hours post-fertilization (hpf) at the 1000-cell stage (15).

In this study, we have combined large-scale transcriptomics and computational analysis to characterize the AS during zebrafish development. Our results show that AS can be divided into two main classes based on their expression correlation with the overlapping protein-coding genes. Surprisingly, these two classes show distinct characteristics in terms of time and space dependent expression, positioning vis-à-vis protein-coding genes, conservation, sequence similarity and transcriptional regulation. These distinct characteristics can be utilized not only in elucidating the role of antisense lncRNAs in development but also in predicting their relation to sense strand gene expression in other biological scenarios.

## MATERIAL AND METHODS

### Identification & Categorization of protein-coding and antisense pairs

To identify and characterize overlapping protein-coding mRNA and antisense non-coding RNA pairs we considered 49672 protein-coding and 1538 antisense transcripts which are annotated in zebrafish genome by ENSEMBL (16). Using the genomic coordinates, we identified all the pairs where the antisense gene coordinates overlap at least 10% of protein-coding gene (1482 pairs). For this, we used BedTools program suit and calculated overlap between pairs using -S option (17).

### Analysis of RNA-Sequencing dataset

FASTQ files corresponding to raw RNA-sequencing reads for the study on 8 zebrafish developmental stages was downloaded from SRA database (https://www.ncbi.nlm.nih.gov/sra). They were quality checked and trimmed using FastQC (developed by Andrew, S., Barbraham Bioinformatics) and Trimmomatic (v0.27) (18), respectively. The reads, for 8 different developmental stages, were then mapped back to the Zv9 or danRer7 genome assembly of zebrafish using TopHat (Bowtie2 v2.1.1) (19). Aligned reads were then used for transcript assembly and expression levels using Cufflinks v2.2.1 (20). For visualisation purposes, strand specific expression tracks were generated using genomecov command in BEDTools package (17).

### Functional analysis of antisense – mRNA pairs

The expression levels of protein-coding and antisense RNA pairs in 8 stages were then used to calculate the correlation between each pair. The pairs were categorized based on whether the pairs correlated positively, negatively or did not show any correlation at all. Only pairs with significant correlation were retained (p-value < 0.05, r >=0.70 for sample size N=8). The stage specific abundance of AS and lincRNAs was visualized using UpSet plots in R software (21). We used Database for Annotation, Visualisation and Integrated Discovery (DAVID 6.8) software for functional annotation of these transcripts (22,23). For this analysis, categories related to biological process, molecular function and cellular components were considered. We used the ggplot2 package in R for plotting all of our violin and box plots, using the geom_violin and geom_boxplot options. We also used the geom_histogram() function to plot the histograms for showing the overlap region and the distance between the TSS of protein-coding genes and the antisense TES. The R software was also used to create heatmaps showing the RNA levels AS and their overlapping protein-coding genes. The table browser from UCSC genome browser was used to download the conservation score (Vertebrate Cons) of each base in the input sequence of AS and mRNA (24).

The G+C content track for zebrafish (danRer7.gc5Base.wig) from UCSC genome browser was used to calculate the G+C content at the promoters (25). We also used EMBOSS geecee analysis to calculate the frequency of G and C nucleotide in the sequences (26).

### ChIP sequencing analysis

The input FASTQ files for published ChIP-Seq data for different histone modifications H3K27me3, H3K4me3 and H3K27ac for Dome (4hpf) stage of zebrafish development were obtained from the DANIOCODE repository (27-29). The ChIP sequencing reads were mapped to Zv9 version of Zebrafish genome using STAR aligner (v2.6.0a) (30). The mapped reads were analyzed using ChIP-seq analysis software HOMER to create tag directories and annotating peaks using the makeTagDirectory and findPeaks tools (-style factor and -o auto options). ChIP-sequencing data was used to plot heatmaps and to visualise the enrichment of different histone modifications within ± 2kb distance of transcription start sites (TSS) of protein-coding transcripts and AS RNAs in three different categories.

### Nuclear & cytosolic fractionation of zebrafish embryos in different stages of development

Wild-type male and female zebrafish (AB-strain) were set up in breeding tanks overnight and the eggs were collected the next day as soon as (∼10 min) they were laid (important to get synchronised embryos). About 100-500 (depending upon the stage) embryos were collected for each developmental stage (32 cells, 64 cells, 256 cells, 512 cells, high, shield, 24hpf, 50hpf). The embryos were treated with Pronase (protease from Streptomyces griseus, Sigma-Aldrich) to remove the chorion, 1mL of Pronase working solution (1mg/mL) was added to 2-3 ml of fish water containing embryos. Dechorionated embryos were then transferred to a 1.5mL microcentrifuge tube and then washed twice with 1 X PBS (Phosphate buffer solution, ThermoFisher). RLN buffer (Sigma) was then added to the tube containing embryos and the yolk sac disrupted releasing the cells buffer. This was then incubated on ice for 5 mins and centrifuged. The resulting supernatant was collected and labelled as the cytosolic fraction. The pellet was washed twice with 200 µL of RLN buffer and finally collected and labelled as the nuclear fraction. RNA was extracted from the nuclear and cytosolic fractions of the embryos in different stages as explained before using the RNeasy mini kit from QIAGEN and DNase (Sigma) treated. The quality of the RNAs extracted was detected on the RNA TapeStation using Agilent High Sensitivity RNA Screen Tape assay. Sequencing libraries were prepared using the TruSeq Stranded Total RNA library Prep kit with the Illumina Ribo-Zero rRNA removal kit (Human-Mouse-Rat). The quality check, library preparation and sequencing were carried out by the University of Birmingham Genomics facility using Illumina NextSeq 500.

The RNA sequencing reads obtained were mapped to the Zv9 genome assembly of zebrafish using STAR (v2.6.0a) (30). Aligned reads were assembled into transcripts using StringTie (31). Further Ballgown program (32) was used to produce differential expression values (in transcript per million or TPM) in different stages of development. These expression values were used for all our downstream analysis.

### CAGE sequencing & analysis

CAGE library preparation was carried out using a modified cap trapping protocol (33) for low quantity samples or LQ-ssCAGE protocol (34). RNA was extracted using the RNeasy mini kit from QIAGEN with 1µg of RNA per sample as starting material. We divided each sample into four parts, 250ng of RNA to be pooled later after cDNA synthesis.

Raw tags from CAGE sequencing were mapped using STAR aligner and the resulting BAM files were used in the bioconductor package CAGEr for further downstream analysis. CAGEr starts from mapped reads and does quality filtering, normalization, removal of the additional 5’ end G nucleotide (added during the CAGE protocol) and the frequency of the usage of start sites (35).

## RESULTS

### Comparison of long intergenic and AS during early development

Understanding the expression patterns of the lncRNAs can give an idea about their role during early zebrafish development. Therefore, we first analyzed expression levels of lncRNAs in the 8 stages of early development. We mined expression data from eight stages of zebrafish development i.e. 2-4 cell, 1000 cell, dome, shield, bud, 24hpf, 48hpf and 120hpf. RNA-seq reads from previously published study (8) was remapped to Zebrafish genome and normalized abundance of annotated lincRNAs, AS and protein-coding genes was calculated. According to ENSEMBL annotations, Zebrafish genome (Zv9, zebrafish release 79) expresses 1538 AS, 4280 long intergenic RNAs (lincRNAs) and 49672 protein-coding (mRNA) transcripts (16). As a first step, we compared the characteristics of the two categories of lncRNAs, *viz*, AS which overlap protein-coding genes and lincRNAs, which are expressed from intergenic regions i.e. away from protein-coding regions of genome (Figure 1). Interestingly, the expression patterns of AS and lincRNAs were distinct. The percentage of 1538 AS present in maternal stages (2-4 cell and 1000 cell) is ∼14% higher than percentage of lincRNAs (Figure 1A). However, after zygotic genome activation, the percentage of AS and lincRNA present is very similar. This indicates that a higher number of AS is deposited among maternal RNAs than lincRNAs suggesting possible relevance of AS in the pre-MZT stages. We also checked the average abundance of AS and lincRNAs. The average level of AS was rather similar or albeit lower as compared to lincRNAs in all the eight stages considered in the study (Figure 1B). This suggests that although the number of AS species present in the pre-MZT is higher than lincRNAs, they are equally abundant to lincRNAs. It is possible that the higher percentage of AS during maternal stages might result from differences in stability in the two types of lncRNAs. Therefore, we also compared the stability and stage specificity of these two classes of lncRNAs. We utilized UpSet diagrams (21) to visualize the frequency of AS and lincRNAs that are present in consecutive stages of development (Figure 1C). We found that the percentage of AS candidates that are present in more than six post-MZT stages is almost half (11.6%) that of the lincRNAs (21%). On the other hand, the percentage of AS RNAs (∼10%) stable during first four stages covering MZT is higher than lincRNAs (∼ 6%). In general, the combined frequency of AS (1.5%) present or expressed in stage specific manner i.e. occurring only in one particular stage was very similar to the lincRNAs (1.1%) (Figure 1C). This suggests that maternally deposited AS RNAs are much more stable as compared to maternally deposited lincRNAs.

**Figure 1:**
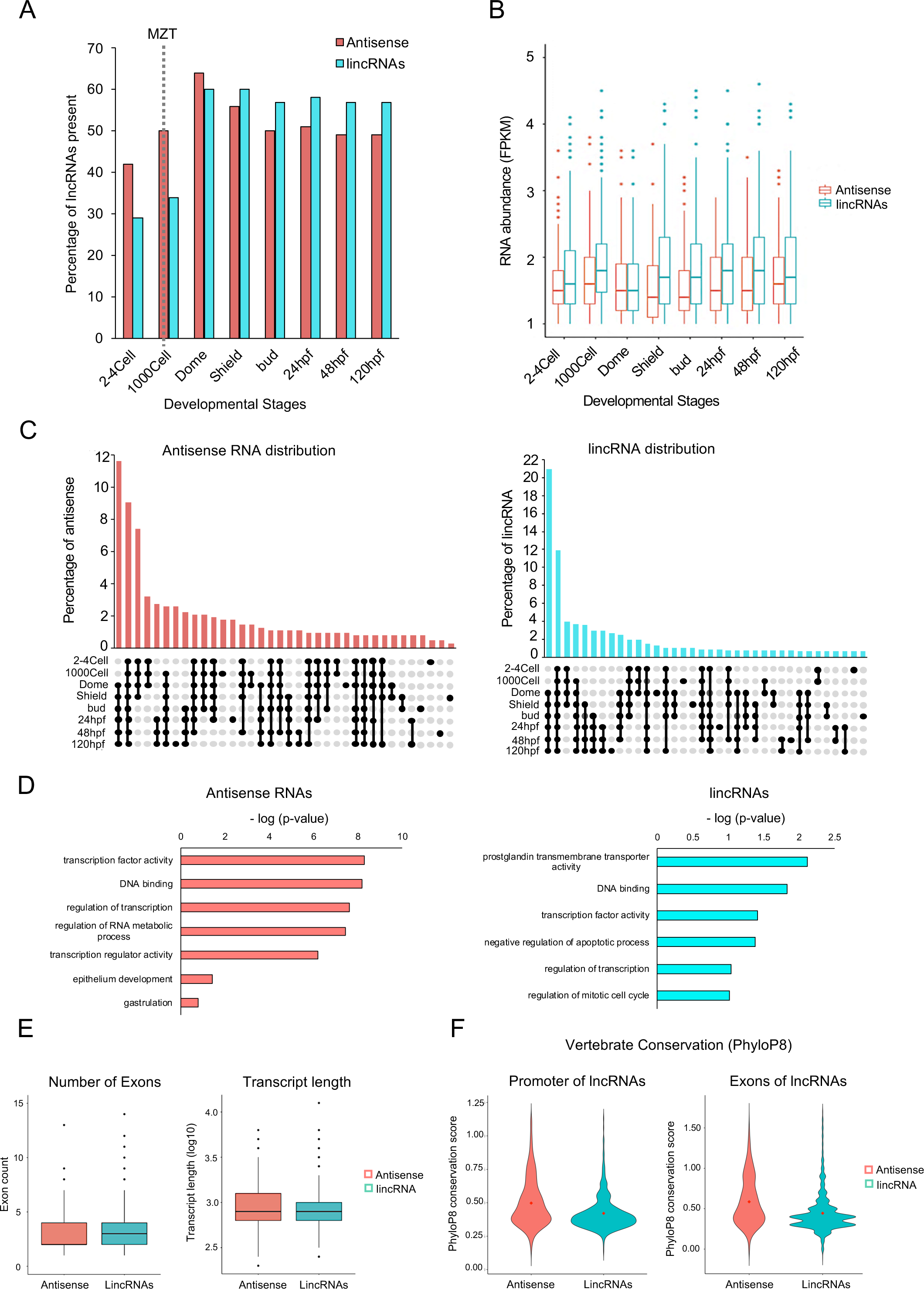
LncRNA dynamics during zebrafish development. **(A)** A bar plot showing the percentage of antisense and lincRNAs present during 8 stages of zebrafish development **(B)** A boxplot of abundance levels of lncRNAs across 8 developmental stages **(C)** UpSet diagrams depicting the percentage of antisense (left) and lincRNAs (right) that are common or unique in the 8 stages of zebrafish development. The percentage of lncRNAs is shown on y-axis and the stages in which the lncRNA is present is shown below x-axis. **(D)** Bar plots of top gene ontology terms associated with the mRNAs that overlap AS (left) vs mRNAs neighboring to lincRNAs (right). The -log(p-val) is plotted on the x-axis and gene ontology terms are shown on the y-axis **(E)** Boxplots showing differences in the number of exon and transcript length. **(F)** Violin plots displaying the conservation score at promoters and exons of antisense and lincRNAs. The antisense are the more conserved class both at the promoters and exons.

Can these differences in expression patterns be related to the functions of protein-coding genes they are associated with? In order to answer this, we compared functions of the protein-coding genes overlapping AS to protein-coding genes adjacent to lincRNAs (Figure 1D). Surprisingly, we noticed that the two groups of lncRNAs also differed in this aspect. The gene ontology analysis showed that AS mostly overlapped genes coding for DNA binding proteins such as transcription factors (p-value <= 5.40× 10^−9^). On the other hand, the protein-coding genes adjacent to lincRNAs (within ±2kb) were enriched for prostaglandin transmembrane transporter activity and this enrichment was much less significant (p-value <= 7.60×10^−3^) as compared to that seen in case of AS.

In addition, AS RNAs were also distinct to lincRNAs in other properties such as transcript length, exon number and conservation. LincRNAs showed a higher exon count while the AS have a higher transcript length compared to the lincRNAs (Figure 1E) probably suggesting possible role of AS in interference through complementarity with overlapping protein-coding genes. Also, the AS were much more conserved both at the promoters (0.49) and exons (0.6) than the lincRNAs (0.42 and 0.48, respectively), indicating their functional importance in early development (Figure 1F).

### Antisense lncRNAs can be divided into three distinct classes based on their relation to overlapping protein-coding genes

The comparison of AS vs lincRNAs (Figure 1) suggests that AS lncRNAs might be needed in early development. Examples show that AS lncRNAs can regulate the expression of overlapping genes. However, relation to their overlapping protein-coding partner in early development is not interrogated in detail. Previous studies show that AS lncRNAs can both, positively or negatively, regulate the expression of their sense strand partner. However, the properties of AS lncRNAs that decide if they will be positive or negative regulators are not understood. Therefore, we further analyzed properties of AS vis-à-vis their co-expression relationship with the overlapping protein-coding genes. Based on ENSEMBL annotations, there are 1482 protein-coding and AS pairs with a minimum overlap of 10% between the lncRNA and the protein-coding counterpart (Figure 2A). To assess co-expression of a AS and its protein-coding partner, a correlation calculation between their RNA levels across the 8 developmental stages was carried out (8). Out of the 1482 protein-coding and AS pairs, 60 pairs did not express at all in any of the developmental stages observed in the study and so were excluded (Figure 2A). Among remaining 1422 protein-coding and AS pairs, 696 protein-coding and AS pairs were negatively correlated while 580 of them were positively correlated and the rest 146 showed no correlation. The pairs with significant correlations (p-value < 0.05 and r > 0.71) were retained for further analysis. As a result, 127 anti-correlated (group-1) and 326 positively correlated protein-coding and AS pairs (group-2) were shortlisted (Figure 2A). The 146 pairs which did not show any correlation were used as a control set (group-3).

**Figure 2.**
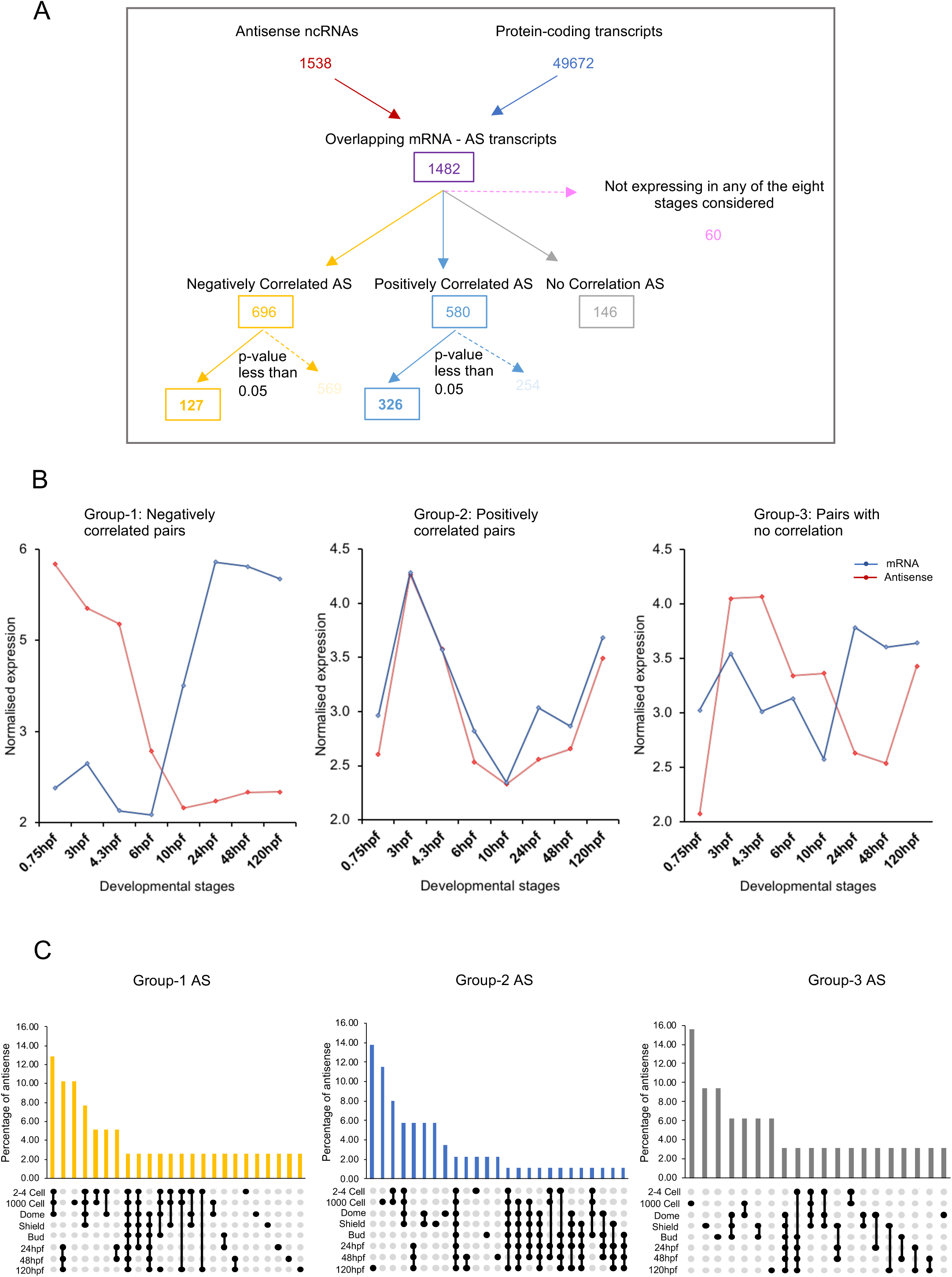
Distinct classes of AS during zebrafish development. **(A)** Overview of the pipeline undertaken to identify and categorize the AS based on their expression correlation with overlapping mRNAs. There are 127 anti-correlated (group-1), 326 positively correlated (group-2) and 146 no correlation (group-3) AS – mRNA pairs. **(B)** Line plots showing the average abundance of AS and overlapping mRNAs in the three categories (negatively correlated, positively correlated and no correlation) during development. **(C)** UpSet plots illustrating the frequency of AS in negatively correlated group-1 (yellow), positively correlated group-2 (blue) and no correlation group-3 (grey) that are common or unique during zebrafish development.

We analyzed the abundance of the AS and protein-coding transcripts in the group-1, group-2 and group-3. In group-1, average abundance of AS was higher in the pre-MZT stages while the protein-coding transcript levels increased post-MZT (Figure 2B, Supplementary figure 1). Thus, suggesting that the AS in the anti-correlated group-1 category were predominantly maternally deposited which eventually get degraded with zygotic genome activation (10hpf). In contrast, in the group-2, both the AS and protein-coding transcripts were present throughout the 8 stages covered by RNA-Seq data (Figure 2B, Supplementary figure 1). Therefore, AS and mRNAs belonging to the positively correlated group showed both maternal (0.75hpf to 10hpf) and zygotic contribution (24hpf onwards). These results were confirmed by plotting individual antisense – mRNA pair RNA levels in the group-1 and group-2 categories in different stages of development (Supplemental Figure 2).

If group-1 is maternally deposited, then RNAs in this group should show distinct stability pre-MZT and post-MZT. In addition, using UpSet plots, we analyzed if AS in the two groups show differences in the stability (Figure 2C). This analysis showed that majority of the antisense in the group-1 category were mostly present either in the early stages (2-4 Cell, 1000 Cell and Dome) or the stages after 24hpf with frequencies 13% and 10%, respectively. The percentage of group-1 AS present only in a single stage was very low suggesting that they are not highly stage-specific. The positively correlated group-2 AS, on the other hand, showed more stage-specific expression with the highest number of AS (abundance > 0.5 FPKM) being specifically present in the 120hpf (14%) and 1000 Cell stage (11%). The control group showed similar stability as in case of group-2 AS. Thus, suggesting that the group-1 AS are more stable and express over a period of time as compared to group-2 AS or the control group. These results show that during early vertebrate development two very different classes of AS are present and they show distinct pattern to their protein-coding partners.

### Developmental genes are associated with AS RNAs in negatively correlated group

Given the distinct expression pattern of genes in the group-1 & group-2, a pertinent question would be if the genes in the positively and negatively correlated group also show differences in their biological and cellular functions. To answer this question, a Gene Ontology (GO) analysis was carried out DAVID program on the protein-coding transcripts in the two groups. The two groups were enriched in distinct molecular functions (Figure 3A). The transcripts belonging to group-1 were enriched in sequence-specific DNA binding proteins (p-value < 3.70 × 10^−9^), developmental genes (p-value 1.40 × 10^−7^) and transcription regulation (p-value < 8.90 × 10^−6^). The top categories in the group-2 showed transcription factor activity (p-value < 1.60 × 10^−4^) although less significant than group-1 mRNAs and housekeeping functions related to metabolism and signaling processes. In contrast, the transcripts in non-correlated group did not show significant enrichment of any particular category.

**Figure 3:**
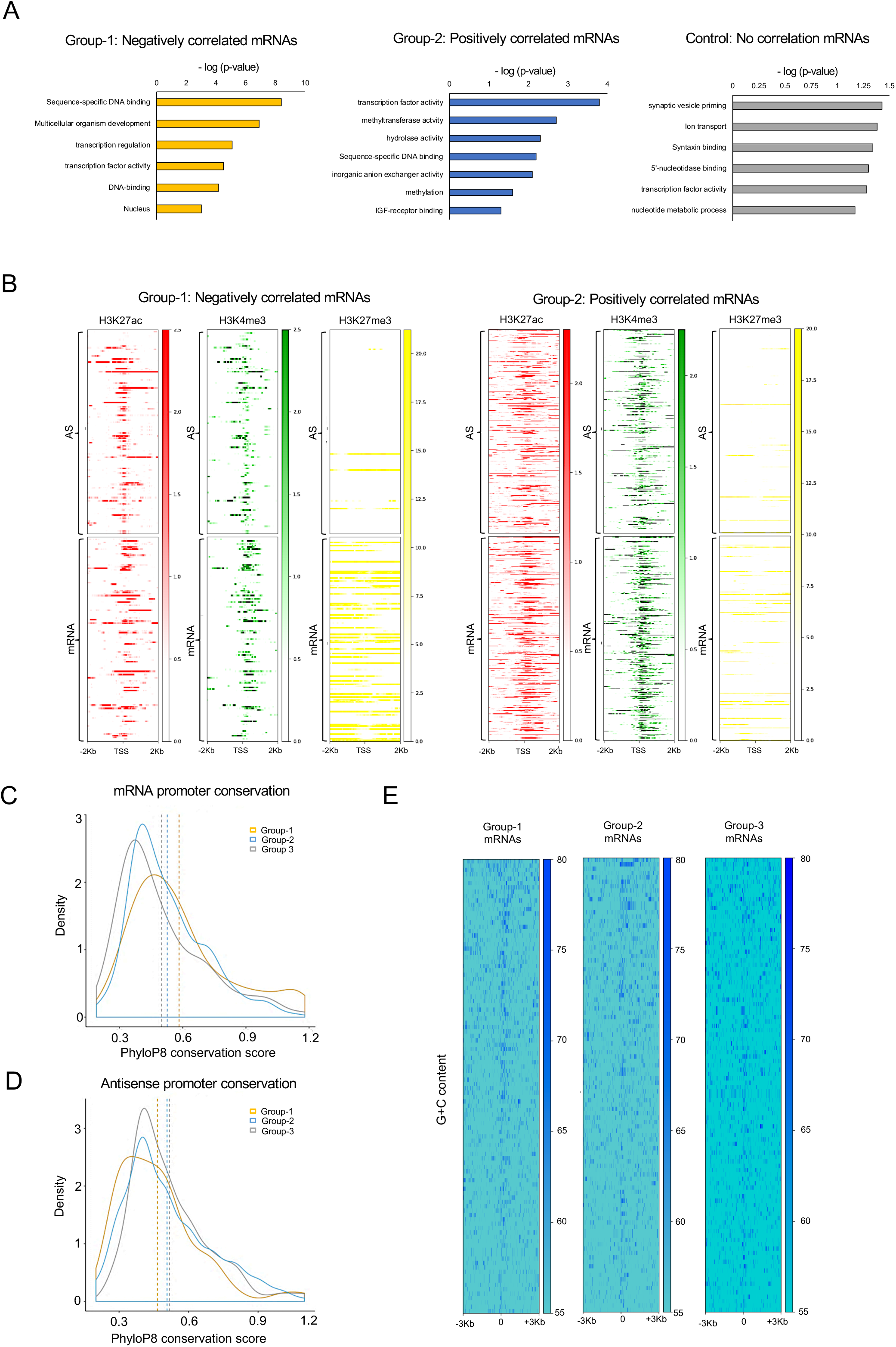
Negatively correlated AS overlap protein-coding genes important in vertebrate development. **(A)** Bar plots depicting the gene ontology terms associated with mRNAs in the three categories AS-protein-coding pairs. In each case the x-axis displays - log(p-value) for each term. **(B)** Heatmaps displaying the distribution of ChIP-Seq reads for H3K27ac (red), H3K4me3 (green) and H3K27me3 (yellow) histone modifications across the TSS of negatively and positively correlated AS (top panels) and overlapping mRNAs (bottom panels). The TSS of mRNAs in the negatively correlated group showed enrichment of H3K27me3 mark suggesting they are targeted by polycomb group of proteins. **(C)** Density plots showing the conservation score (PhyloP8) at the promoters of mRNAs and **(D)** AS in the negatively correlated group-1 (yellow), positively correlated group-2 (blue) and no correlation group-3 (grey). The AS in the negatively correlated group showed a lower conservation score compared to the AS in the other two group. However, the negatively correlated mRNAs are more conserved in comparison. **(E)** Heatmap displaying G+C content at the promoters of mRNAs in the three categories.

It is well-documented that developmental genes are repressed by polycomb group of proteins which catalyze Histone 3 lysine 27 trimethylation or H3K27me3 histone modification marks at the promoter of these genes (36). Given enrichment of developmental genes in group-1 (Figure 3A), we wondered whether the protein-coding genes in this group show enrichment of H3K27me3 or any particular histone modification marks at their promoter during early stages of development. To verify this, we mined genome-wide <u>Ch</u>romatin <u>I</u>mmunoprecipitation data (ChIP-seq) for H3K27me3, H3K4me3 (Histone 3 lysine 4 trimethylation) and H3K27ac (Histone 3 lysine27 acetylation) marks in the dome stage of zebrafish development. H3K27me3 mark is generally enriched at transcriptionally repressed genes while H3K4me3 and H3K27ac marks are usually enriched at transcriptionally active genes. We mapped these marks on Transcription Start Sites (TSS) of protein-coding and AS genes in group-1, group-2 and the control group-3 (Figure 3B). Interestingly, we observed distinct patterns of histone modifications across TSS of antisense and overlapping protein-coding genes in the two groups (Figure 3B). The protein-coding genes (lower panel) in group-1 showed high enrichment of repressive H3K27me3 mark, typically seen at developmental genes. Interestingly, AS in the group-1 showed little or no repressive marks around the TSS. H3K27ac and H3K4me3 marks, which represent activation and initiation marks at the promoter regions, did not show any significant enrichment on either the AS or protein-coding genes of this negatively correlated group. The lack of H3K4me3 and H3K27ac marks and enrichment of H3K27me3 mark supports that AS as well as the overlapping mRNA in group-1 are not transcribed. This is in accordance with our prediction that AS are maternally deposited, and the overlapping mRNAs are not expressed in early stages. In contrast, the histone modification marks, at AS-protein-coding pairs in the group-2, are distinct. The AS as well as the protein-coding genes in the group-2 did not display any enrichment for H3K27me3 repressive mark but there is significant enrichment of activation marks H3K27ac and H3K4me3 across their TSS (Figure 3B right hand panels). This is expected given the genes in positively correlated group are house-keeping genes. The histone modification patterns at group-2 are very similar to average protein-coding genes indicating that group-2 represents housekeeping genes (Supplementary figure 3A). On the other hand, group-3 AS and protein-coding genes did not show enrichment for any histone modifications (Supplementary figure 3B).

Evolutionary conservation of genes or genomic elements is often used as a proxy for their functional importance. We assessed evolutionary conservation of the gene promoters, exons, 3’-UTR and 5’-UTR in the two categories (Figure 3C, D & Supplementary figure 3C & 4A). The protein-coding genes in the group-1 showed higher conservation score at the promoters (0.58) and exons (1.01) compared to the group-2 and the control group-3 (Figure 3C & Supplementary figure 3C). The higher conservation level of protein-coding genes is as expected for developmental genes (Figure 3C). In contrast, however, the promoters (0.46) but not exons of negatively correlated AS showed less conservation than the other two classes of AS (Figure 3D & Supplementary figure 3C). However, the 5’-UTR and 3’-UTRs of both group-1 and group-2 protein-coding genes are less conserved compared to group-3 (Supplementary figure 4A). Since the polycomb group of genes have GC-rich promoters, we further analyzed the GC content at the promoters. Our analysis suggests that the group-1 mRNAs have promoters with higher G+C content as compared to group-2 mRNAs and mRNAs in no correlation group-3 (Figure 3E). The G+C content at 5’-UTR and 3’-UTRs, however, was not different between the protein-coding genes in the three classes (Supplementary figure 4B).

These observations support that the protein-coding genes in group-1 are polycomb targeted developmental genes. On the other hand, group-2 and non-correlated groups were mostly house-keeping genes suggesting a different mechanism of regulation.

### Group-1 AS start in intergenic regions and show greater overlap with protein-coding genes

It has been proposed that the AS regulate expression of overlapping protein-coding genes. However, examples supporting positive as well as negative regulation are described in literature (6). In addition, in many cases, AS did not show any relationship to the overlapping protein-coding gene. This is reflected in our analysis which shows that antisense RNA-protein pairs can be divided into either negative (group-1), positive (group-2) or no correlation groups (group-3). However, it is not clear if these three categories differ in antisense RNA organization with respect to the overlapping protein-coding gene. Understanding how different categories of AS are organized in the genome vis-à-vis overlapping protein-coding genome would help in understanding the mechanisms of AS-mediated regulation of protein-coding gene.

As a first step, we calculated the extent of the overlap between the AS and protein-coding gene pairs in the three categories (Figure 4A). Interestingly, AS-protein-coding genes pairs in group-1 showed much greater overlap (average = 13003bp) as compared to the pairs in group-2 (average = 4548bp) and no correlation group (average = 6722bp). We also analyzed the distances between the annotated (ENSEMBL) transcription start site (TSS) and transcription end site (TES) of AS to the TSS of overlapping mRNA gene (Figure 4B, C). The TSS-TSS (Figure 4B) and TES-TSS (Figure 4C) distance distribution shows that the three groups of AS are quite distinct from one another. In group-1, TSS of AS RNAs is much further away from the TSS of overlapping protein-coding genes (log of distance = 5) as compared to that in group-2 (log of distance = 4). On the other hand, the TES of group-1 and group-2 AS are at similar distance to TSS of protein-coding genes. This is reflected in the greater overlap between the AS and corresponding protein-coding genes in the group-1 category (Figure 4A).

**Figure 4:**
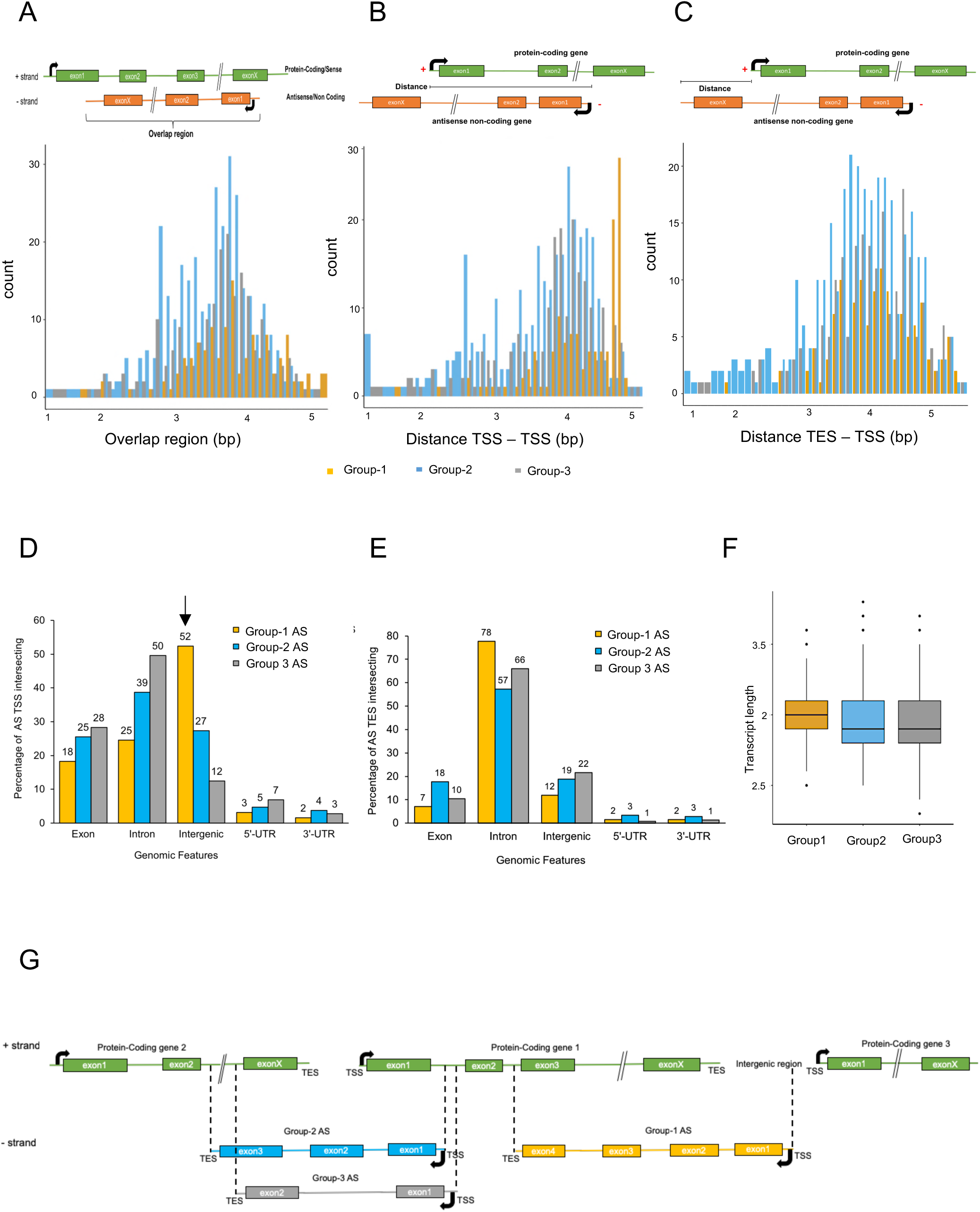
The AS in three categories are differently transcribed vis-à-vis overlapping protein-coding genes. **(A)** A histogram showing the overlap region between the AS and the overlapping mRNAs in the group-1 (yellow), group-2 (blue) and group-3 (grey). **(B)** A histogram showing the distance between the TSS of AS and TSS of overlapping mRNA (log of distance in bp) in the three categories of genes as described in A. **(C)** A histogram showing the distance between the TES of AS and TSS of overlapping mRNA (log) in the three categories. The schematics above each histogram represent how the respective distances were calculated. (**D)** A bar plot of the percentage of AS TSS overlapping different genomic features (exon, intron, intergenic region, 5’-UTR and 3’-UTR) on the opposite strand. **(E)** A bar plot of the percentage of AS TES overlapping different genomic features on the opposite strand. **(F)** A boxplot of log(transcript lengths) of AS in the negatively, positively and no correlation groups. **(G)** Schematic diagram representing how the three different groups of AS are distributed and localized in the genome with respect to their mRNAs.

We further interrogated the location of AS TSS and TES vis-à-vis genomic features such as exon, intron, intergenic, 3’-UTR and 5’-UTR of protein-coding genes on the opposite strand (Figure 4D, E). Remarkably, majority of AS in group-1 started in intergenic regions in contrast to the group-2 and no correlation groups where AS RNAs mainly started within the protein-coding gene overlapping opposite strand (Figure 4D). On the contrary, the TES of group-1 AS overlap the introns on opposite strand which is also true for the AS in the other two categories of AS (Figure 4E). Interestingly, the larger overlap between AS and protein-coding genes in group-1 is also reflected in the larger transcript-length of AS in group-1 as compared to group-2 and non-correlated group (Figure 4F). The number of exons was also higher for group-1 AS (Supplementary figure 5A). These observations together show that AS in group-1 start in intergenic regions neighboring the TES of overlapping protein-coding gene. Group-1 AS being longer span most of the length of overlapping protein-coding gene and end nearer to TSS of the protein-coding gene (Figure 4G). Group-2 AS, on the other hand, start in exonic or intronic region nearer to the TSS of overlapping protein-partner. They end near the TSS of overlapping protein-coding gene pair (Figure 4G).

### AS in the two categories show different transcription regulation

The distinct expression patterns of AS and different functional roles of overlapping protein-coding genes in the three different categories point to possible differences in transcription regulation of AS. This is also supported by distinct locations of TSS of AS RNA in these categories (Figure 4D). To further dissect the mechanism of their regulation, we analyzed sequence motifs associated with the promoters of AS in these three categories. Three categories showed enrichment of distinct transcription factor (TF) motifs (Figure 5A). We observed that the AS in the negatively correlated group-1 showed enrichment for TFs such as MTF1 and RUNX3 which are shown to be essential for normal vertebrate development (37-39). The group-2 category AS, on the other hand, showed enrichment for HES7 and IRF-6 transcription factors (Figure 5A) also involved in zebrafish development.

**Figure 5.**
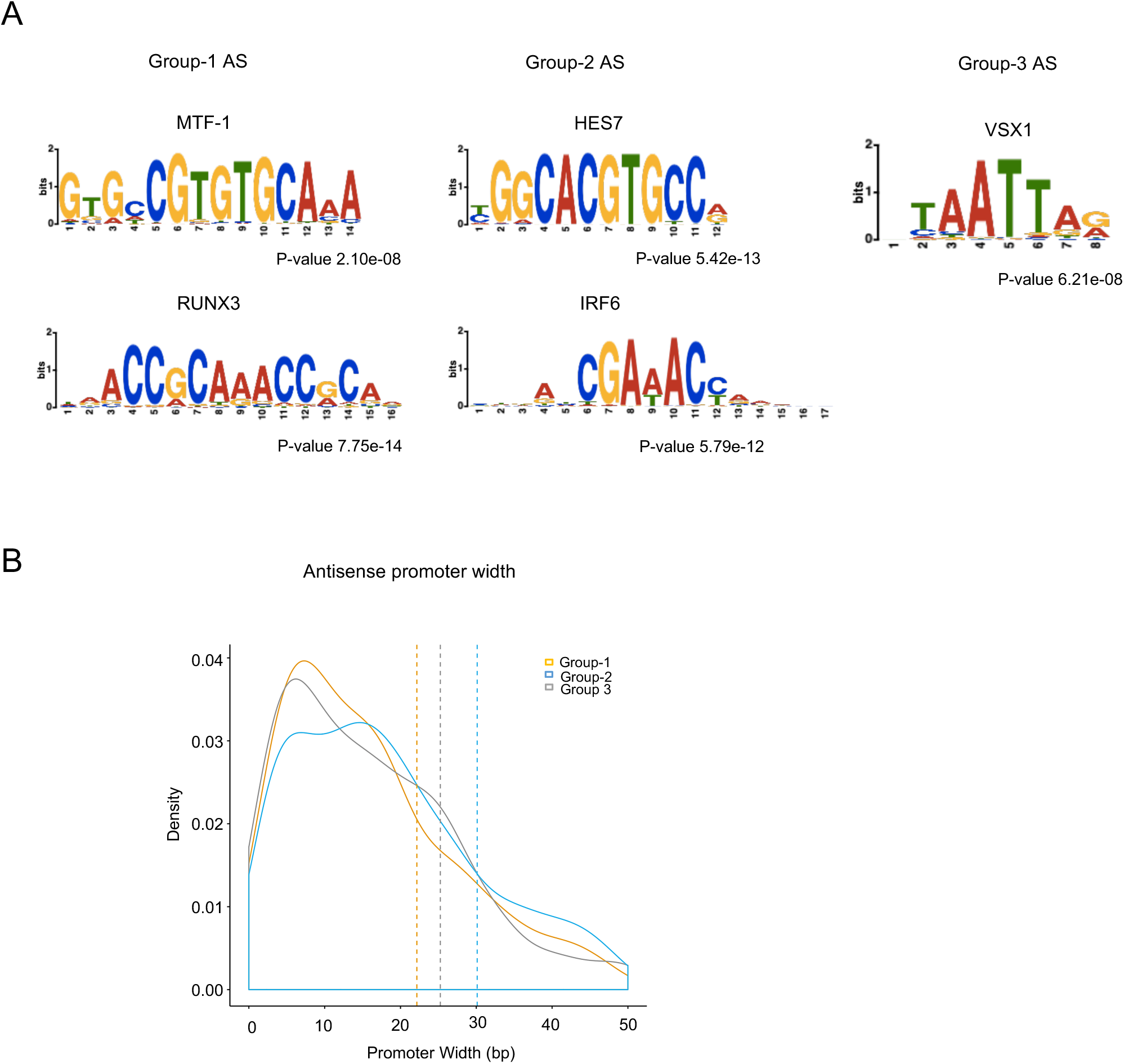
Analysis of motifs associated with the AS and mRNAs in the three different categories. **(A)** the figure shows the different motifs associated with the TSS of AS and overlapping mRNAs in the negatively, positively and no correlation group. **(B)** graph for promoter-width (x-axis) distribution of the AS in the three groups.

Another indication of transcription regulation can be provided by promoter width. It has been reported before that tissue-specific promoters tend to be sharper than house-keeping genes which tend to have broader promoter (35). The promoter width can be effectively calculated by mapping TSS using <u>C</u>ap <u>A</u>nalysis <u>G</u>ene <u>E</u>xpression analysis or CAGE analysis. CAGE is a method for transcriptome analysis about changes in TSS and its relative usage at single nucleotide resolution (40). CAGE gives us the information for the start sites of capped RNAs which in turn can be used as an indicator of promoter organization. The width of TSSs are calculated using the mapped CAGE tags. For this analysis, we deep sequenced the CAGE tags from six stages of early zebrafish development and calculated the promoter width (Figure 5B). In this analysis, promoters of positively correlated Group-2 AS RNAs appeared distinct as compared to Group-1 and no-correlated control group. This observation again supports our previous analysis that Group-1 AS are more tissue-specific as compared to Group-2 which show promoters with characteristics similar to house-keeping genes.

### AS in different categories show differences in cellular localization

Studies on yeast (41) and flies (42) have indicated very specific subcellular localization of mRNAs to be important in yeast and fly development. It can be envisioned that the specific subcellular localization of AS during zebrafish embryogenesis might also be essential for their regulatory function. Therefore, we sought to identify the subcellular localization of RNAs using total RNA-seq (both polyA+ and polyA-) on nuclear and cytosolic fractions from early stages of zebrafish embryos. This can not only provide information on the localization of developmental mRNAs, but also several processed and unprocessed non-coding RNAs. The fractionation of RNAs will also allow detection of lowly expressed RNAs which cannot be detected in whole-cell RNA-seq due to low transcript levels and being restricted to the nucleus. To define an RNA as nuclear or cytosolic, we used the log2 ratio of nuclear and cytosolic RNA levels (Nuclear/Cytosolic). An RNA was categorized as nuclear, if the ratio was greater than 0.65 (1.5-fold enrichment), and cytosolic, if it was less than -0.65 (1.5-fold enrichment). And anything in between the ratios (0.65 and -0.65) was considered to be more or less equally present in both cellular fractions.

First, we analyzed the localization and expression of annotated mRNAs and lncRNAs in the nuclear and cytosolic compartment during zebrafish development. In general, we found that AS primarily dominate in the cytosolic fraction while lincRNAs are mostly nuclear in the pre-MZT stages (Figure 6A). However, with zygotic genome activation lincRNAs show similar abundance in nuclear and cytosolic fractions of the cell.

**Figure 6.**
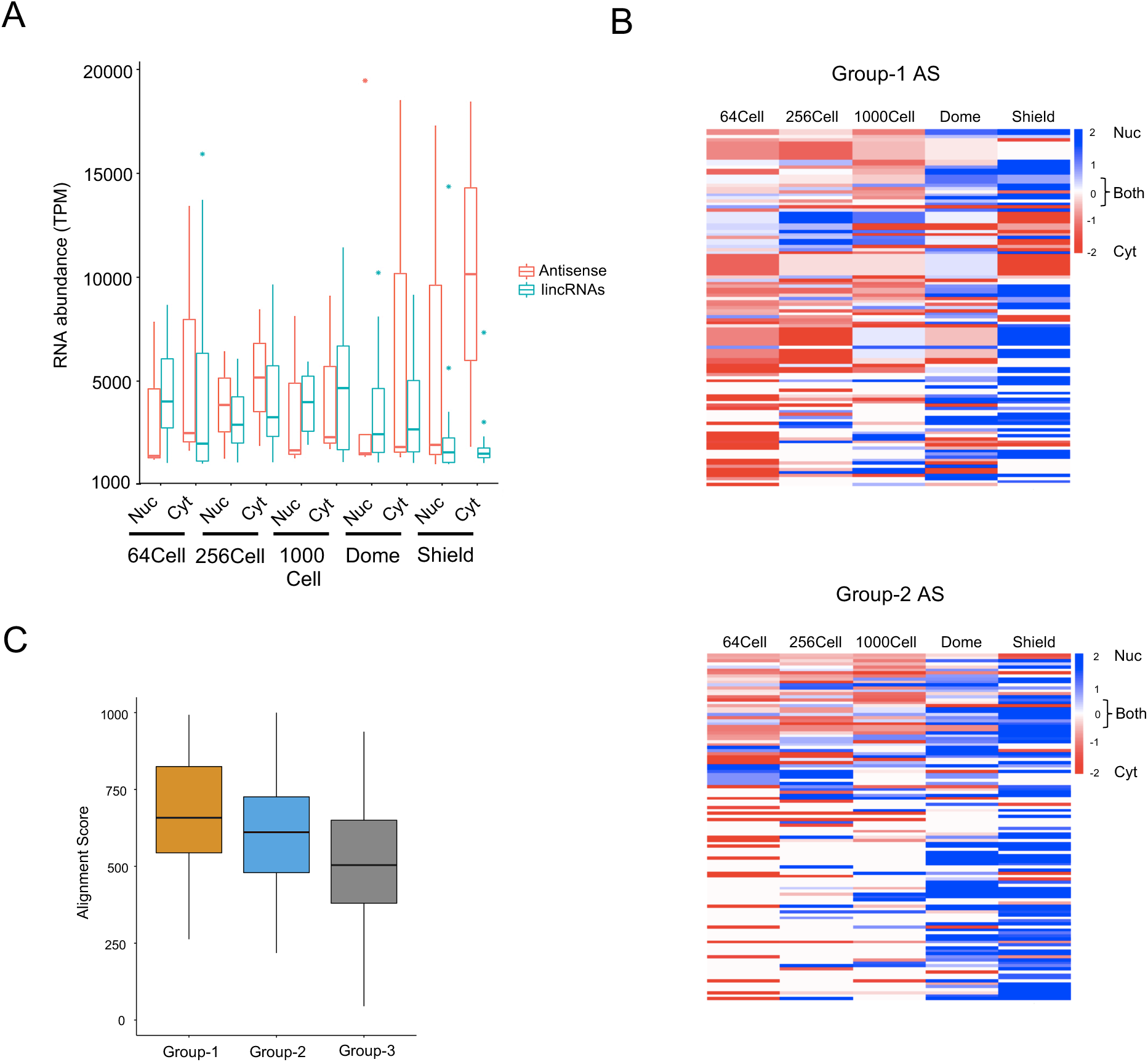
Localization of lncRNAs with zebrafish embryogenesis. **(A)** A boxplot comparing the percentage of antisense and lincRNAs that are nuclear or cytosolic. **(B)** Heatmaps showing the localization of group-1 and group-2 AS during zebrafish development. The scales are from -2 (red, cytosolic) to 2 (blue, nuclear) and 0 (white, both). **(C)** A boxplot of the alignment score between AS and mRNA sequence in the group-1 (yellow), group-2 (blue) and group-3 (grey) group.

In addition, we also analyzed if AS in different categories show differences in the cellular localization and if this can give us a clue regarding their function. We ranked the AS based on their abundance and then calculated the log2 ratio to determine the localization of these AS. The group-1 AS were more enriched in the cytosolic fraction during pre-MZT stages further supporting maternal deposition and become more enriched in the nuclear fraction in the shield stage when zygotic transcription begins (Figure 6B above). Majority of group-2 AS are either present in both the fractions and are enriched in the nucleus in the stages after 64 cell stage (Figure 6B below). The control group did not show any particular enrichment (Supplementary figure 5B).

Previous studies have indicated that cytosolic AS downregulate their overlapping protein-coding gene function by forming an RNA-RNA hybridization with the mRNA thus preventing its translation (43). A similar functional mechanism can be envisioned for group-1 AS given that they are cytosolic and show negative correlation with expression levels of their protein-coding partners. Therefore, we assessed the sequence complementarity between the AS transcript and mRNA in the three groups (Figure 6C). In this analysis, group-1 AS showed a higher alignment score (650bp) for the negatively correlated category compared to the other categories (Figure 6C). This is also reflected by the higher alignment length (1500bp) and a higher complementarity (390bp) between the AS and the mRNA sequence which further validates our theory of post-transcriptional mechanism of regulation of mRNA expression by overlapping antisense (Supplementary figure 5C, D).

## DISCUSSION

Antisense transcription is a common feature among all types of organisms ranging from bacteria to mammals with up to 20% of human transcriptome being predicted to show evidence of antisense transcription (3,44). Examples show that antisense RNAs regulate the overlapping protein-coding gene on sense strand. It has become clear that a large proportion of protein-coding genes are part of sense-antisense transcript pairs where the antisense transcript in many cases is a lncRNA (44-49).

Individual examples indicate that the antisense ncRNAs employ various mechanisms to regulate overlapping mRNA expression. Some regulate transcription by influencing epigenetic modifications (50) or recruitment of transcription factors (51,52). Other examples suggest their role in regulating alternative splicing and translation (53-56). Recent studies also show that antisense RNAs are the most predominant ncRNAs in early development (8). However, few of them are studied in detail and are shown to play a role in early development (7). Unsurprisingly, number of antisense RNAs are implicated in many diseases (47,57-59).

Despite the widespread nature of antisense transcription, they remain poorly characterized. Their functional relation to the overlapping sense strand protein-coding genes remains confusing. One of the reasons behind this is that antisense RNAs display positive as well as negative correlation to the expression of their sense strand partners. In addition, a number of antisense RNAs do not affect the expression of overlapping genes. As a result, it has been difficult to propose unifying mechanism of their function. Instead, we took a different approach where we first divided antisense RNAs based on their co-expression patterns vis-à-vis their overlapping protein-coding partners and then studied the features of antisense RNAs in each group. We focused on analyzing characteristics of the two sets of antisense ncRNAs across eight stages of zebrafish development i.e. with negative (group-1) and positive (group-2) co-expression patterns with overlapping protein-coding genes as compared to those which do not show any relation to the overlapping protein-coding gene (group-3).

Surprisingly, the two sets of antisense RNAs and protein-coding genes show very distinctive characteristics. Group-1 protein-coding genes are mainly developmental genes while Group-2 protein-coding genes show enrichment for house-keeping functions. On the other hand, Group-3 did not show enrichment for any particular category of proteins. Interestingly, Group-1 and Group-2 antisense RNAs were also distinctive in terms of transcript lengths, promoters, transcription start sites and transcription end sites, indicating that their expression is regulated in different manner. This is reflected in their expression pattern during development with Group-1 antisense RNAs being mainly maternally deposited while Group-2 RNAs seem to have transcribed from zygotic genome. Interestingly, our RNA-seq data shows that Group-1 RNAs are cytosolic which is typical of maternal RNAs. Group-2 RNAs on the other hand show nuclear presence, possibly because they are transcribed from zygotic genome.

The studies on antisense RNAs have shown they can regulate either transcription or post-transcriptional processing. NcRNAs which regulate transcription are generally nuclear. Majority of them act by changing chromatin landscapes around protein-coding genes. The antisense RNAs which act post-transcriptionally generally act by changing stability of complimentary mRNAs, affecting splicing pattern or the rate of translation (60).

Given cytosolic nature of Group-1 RNAs and their opposite expression pattern as compared to developmental mRNAs, we can speculate that they might be involved in downregulating expression of developmental mRNAs during early stages of zebrafish development. They might be involved in decreasing mRNA stability through microRNA-like mechanism of hybridization-mediated degradation. Higher level of complementarity in the Group-1 antisense RNA-mRNA pairs as compared to other two groups corroborates possible hybridization between antisense RNA and mRNA. However, why do we need downregulation of developmental mRNAs which are transcribed only after MZT? And in that case what is the reason behind the higher percentage of Group-2 antisense RNAs among maternally deposited RNAs? It is possible that they do not have any significant role in regulating the developmental genes. However, their expression specifically from developmental gene loci and the distinct features is not trivial. Another explanation is that unwarranted early expression of developmental genes can be detrimental to normal development. Therefore, we speculate that these RNAs might form a second level of regulation to downregulate any precarious expression of developmental genes. This hypothesis however needs further experimental verification.

In addition to functional differences between Group-1 and Group-2 mRNAs, the differences in their genomic configuration are interesting. Group-1 ncRNAs start in intergenic region; away from TSS of their overlapping protein-coding partner and generally display high overlap with the sense gene. They tend to be in head-to-tail configuration w.r.t. sense gene. In contrast, Group-2 ncRNA TSSs are much closer to sense gene TSS and they appear to have more head-to-head configuration vis-à-vis sense protein-coding gene. In future, these features can be used to predict relationship between antisense ncRNAs to their overlapping protein-coding genes. An in-depth study is however needed to further understand the relationship antisense ncRNAs and their co-expression pattern with their protein-coding partner.

## AVAILABILITY

https://bedtools.readthedocs.io/en/latest/content/overview.html

https://www.bioinformatics.babraham.ac.uk/projects/fastqc/

http://www.usadellab.org/cms/?page=trimmomatic

https://ccb.jhu.edu/software/tophat/index.shtml

http://cole-trapnell-lab.github.io/cufflinks/

https://danio-code.zfin.org/

https://cran.r-project.org/web/packages/UpSetR/vignettes/basic.usage.html

https://david-d.ncifcrf.gov/

https://www.rdocumentation.org/packages/ggplot2/versions/3.2.1

https://genome.ucsc.edu/cgi-bin/hgTables

http://www.bioinformatics.nl/cgi-bin/emboss/geecee

http://labshare.cshl.edu/shares/gingeraslab/www-data/dobin/STAR/STAR.posix/doc/STARmanual.pdf

http://homer.ucsd.edu/homer/ngs/index.html

https://ccb.jhu.edu/software/stringtie/

GEO/GSE32898

GEO/GSE32899

GEO/GSE32900

## ACCESSION NUMBERS

The RNA-seq and CAGE-seq data generated in this study is deposited in GEO database under accession numbers GSE143208 and GSE144040 respectively.

## SUPPLEMENTARY DATA

Supplementary Data are available online.

## ACKNOWLEDGEMENT

We thank Prof Ferenc Mueller and his lab for useful discussions and help with zebrafish methods. We are grateful to all members of ZENCODE ITN for critical comments on the work. We also thank Nishiyori Hiromi and Miki Kojima from Laboratory for Transcriptome Technology, RIKEN Center for Integrative Medical Sciences, Japan for their help with LQ-CAGE library preparation.

## FUNDING

This work was supported by the European Commission’s Marie Curie H2020 ITN funding. Funding for open access charge: University of Liverpool.

## CONFLICT OF INTEREST

Authors declare no conflict of interests.

## Notes

#### Summary of Updates

Minor formatting corrections

