## Supplementary figures for "Antisense ncRNAs during early vertebrate development are divided in groups with distinct features"

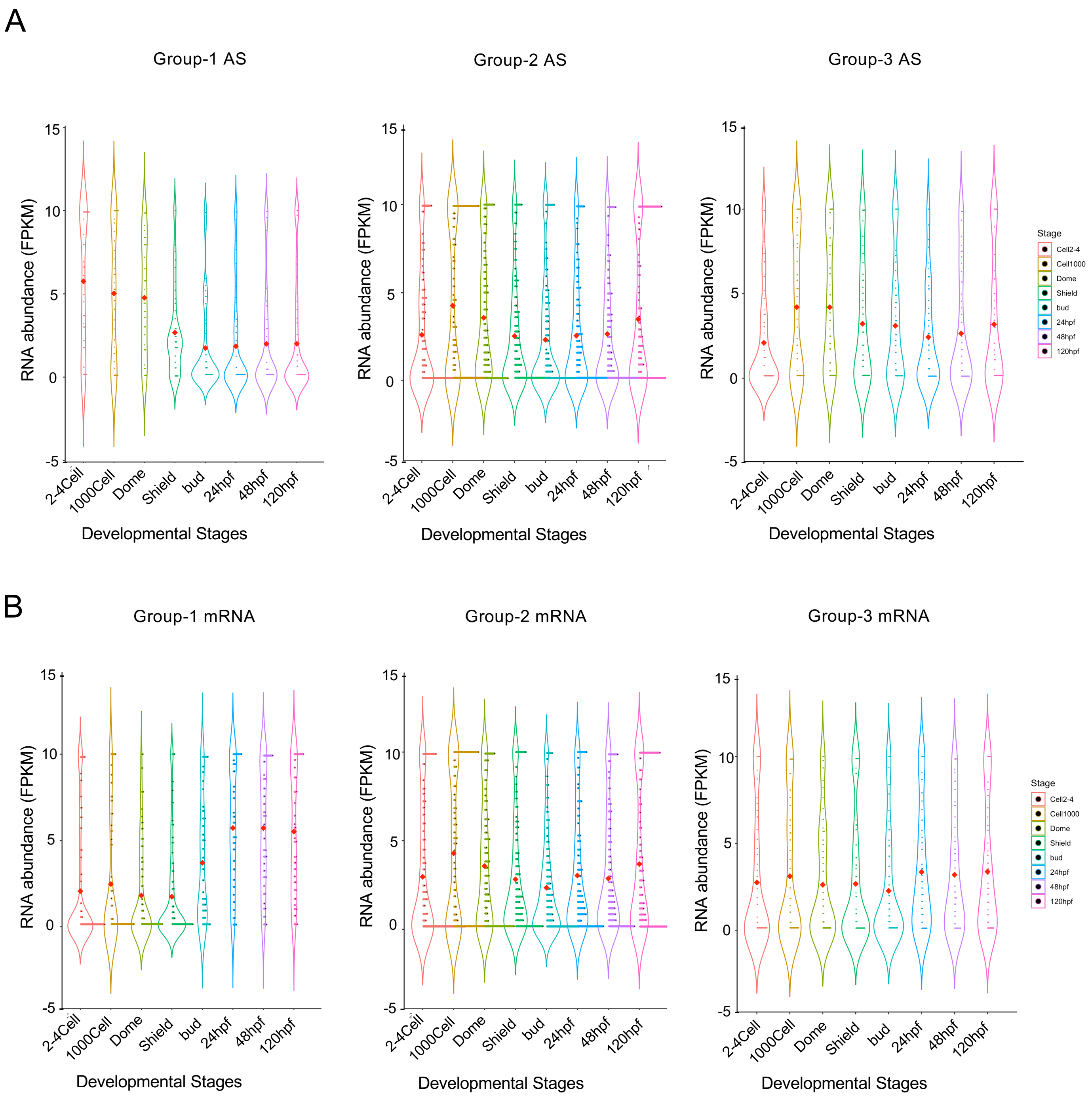


Supplementary Figure 1. (A) Violin plots showing the levels of antisense RNA in group-1, group-2 and group-3 categories. (B) Violin plots showing the levels of overlapping mRNAs in group-1, group-2 and group-3 categories. The Y-axis represents the abundance of RNAs (FPKM) while the X-axis shows the stages of development.

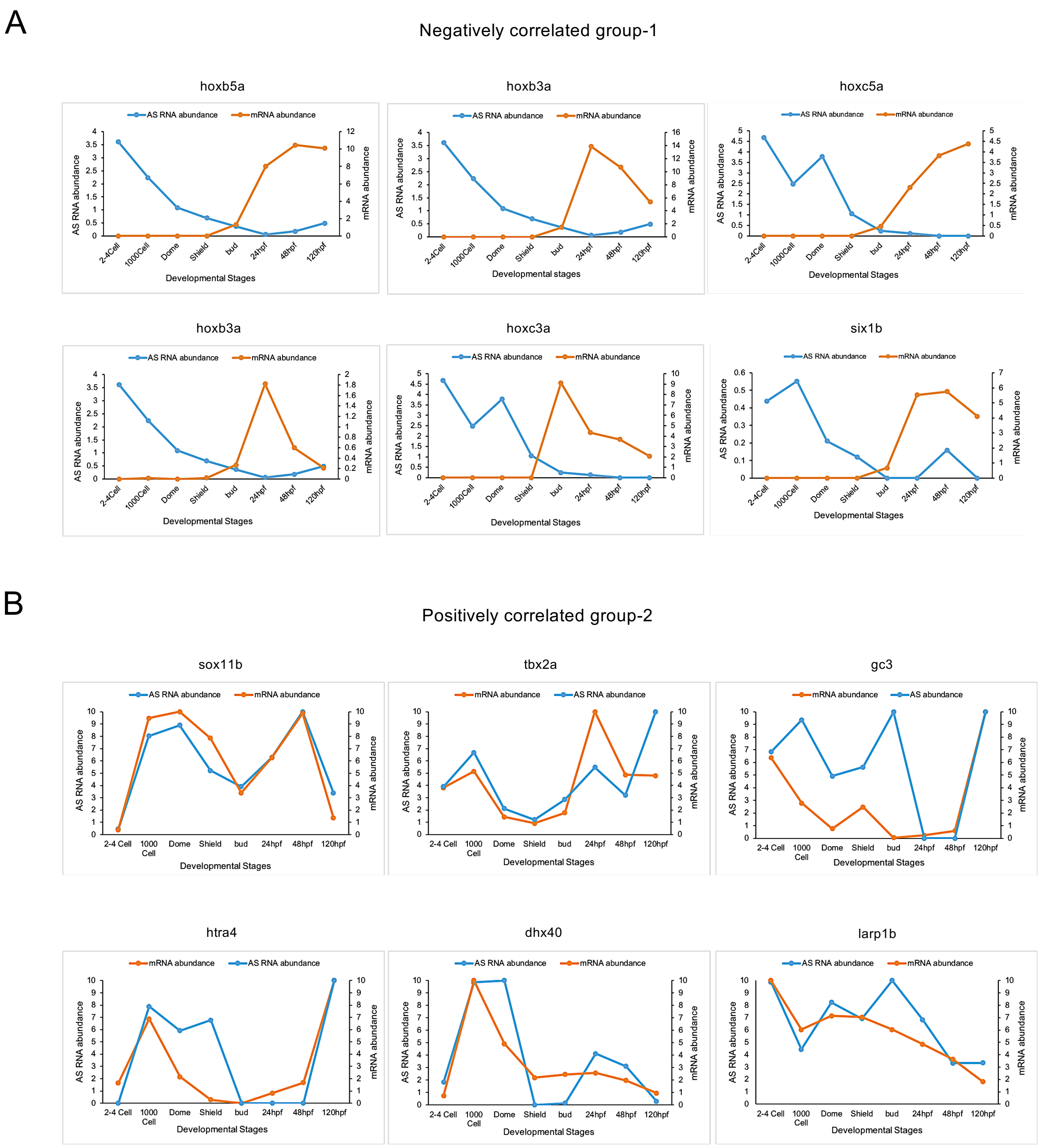


Supplementary Figure 2. (A) Individual examples showing levels of antisense RNA and overlapping mRNA in the group-1 category. (B) Individual examples showing average RNA levels of antisense and mRNAs in group-2 category. The Y-axis represents the abundance of RNAs (FPKM) while the X-axis shows the stages of development.

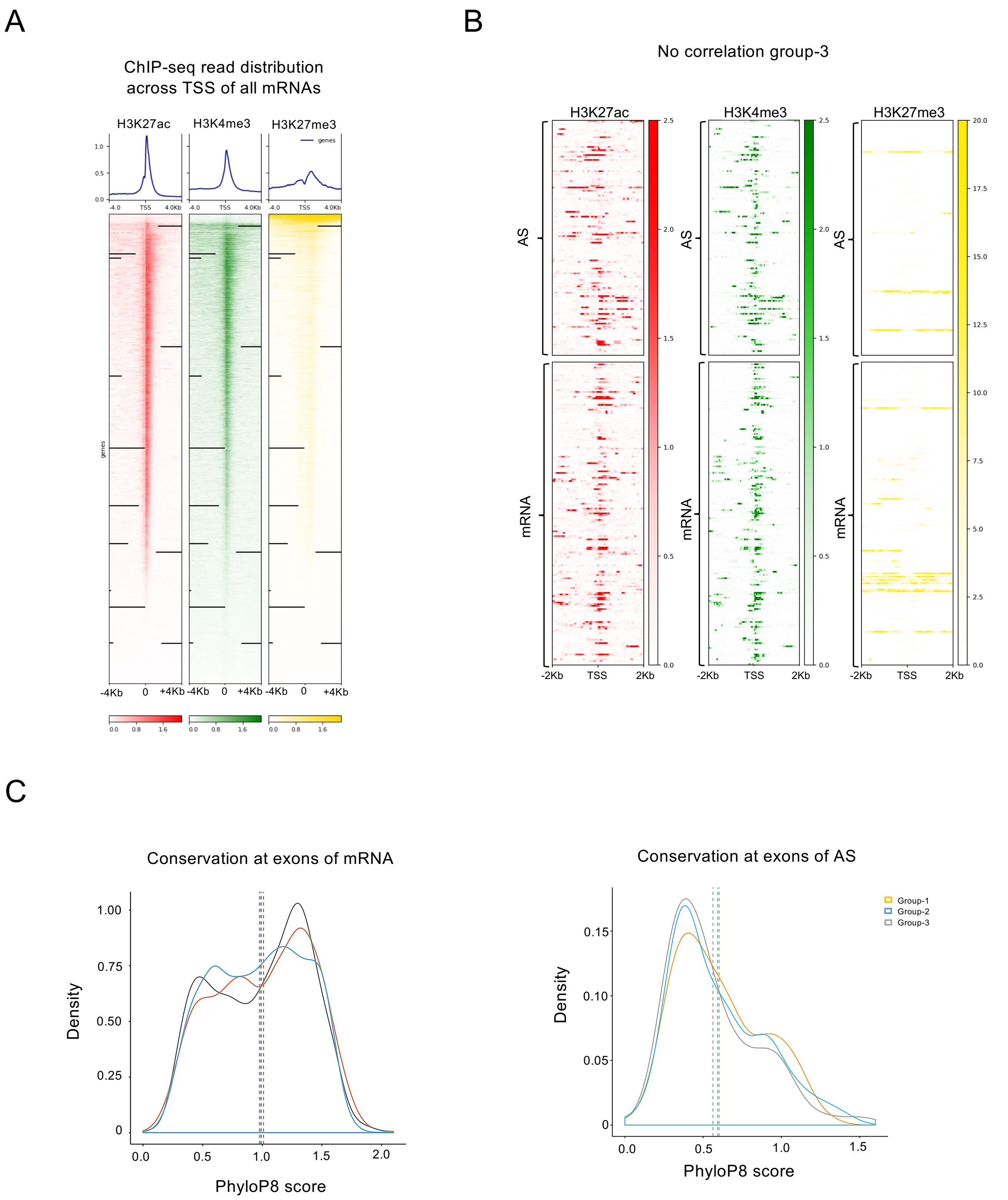


Supplementary Figure 3. (A) Heatmaps showing the enrichment of H3K27ac (red), H3K4me3 (green) and H3K27me3 (yellow) ChIP-seq reads across the TSSs of annotated mRNAs (±4kb). A very clear enrichment is seen for all the modifications considered in the study. (B) Heatmaps showing the distribution of H3K27ac (red), H3K4me3 (green) and H3K27me3 (yellow) ChIP tags across the TSSs of Group-3 AS and mRNAs. (C) Density plots showing the conservation at exons of protein-coding genes (left) and antisense RNA genes (right) in the three groups.

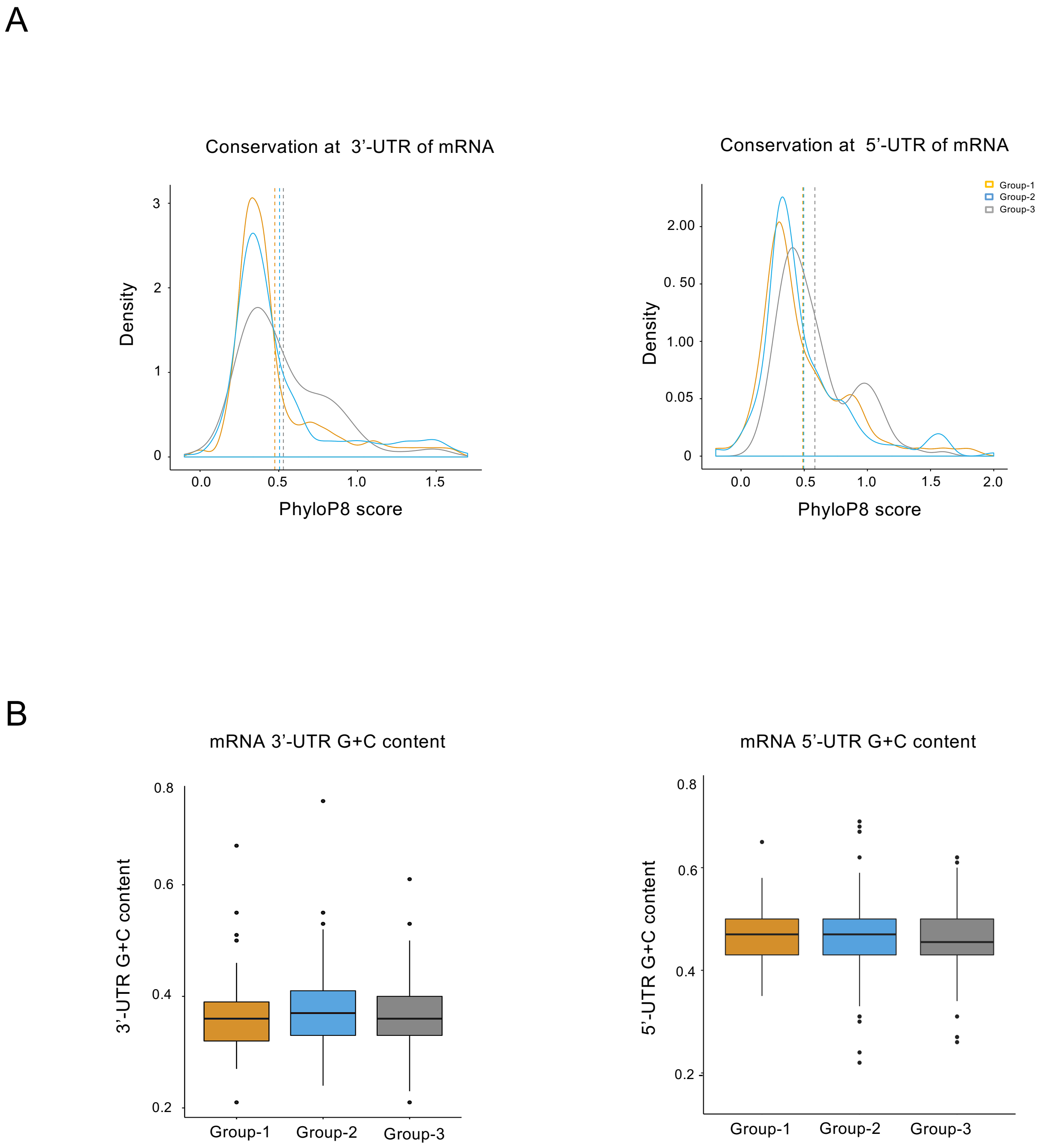


Supplementary Figure 4. (A) Density plots of the conservation at 3'-UTR (left) and 5'-UTR (right) of protein-coding genes in the three groups. Both the group-1 & group-2 mRNAs are less conserved at 3'-UTRs but more conserved in the 5'-UTRs compared to the group-3. (B) Boxplots of G+C content at the 3'-UTR (left) and 5'-UTR (right) of group-1, group-2 and group-3 mRNAs using EMBOSS-6.6.0.

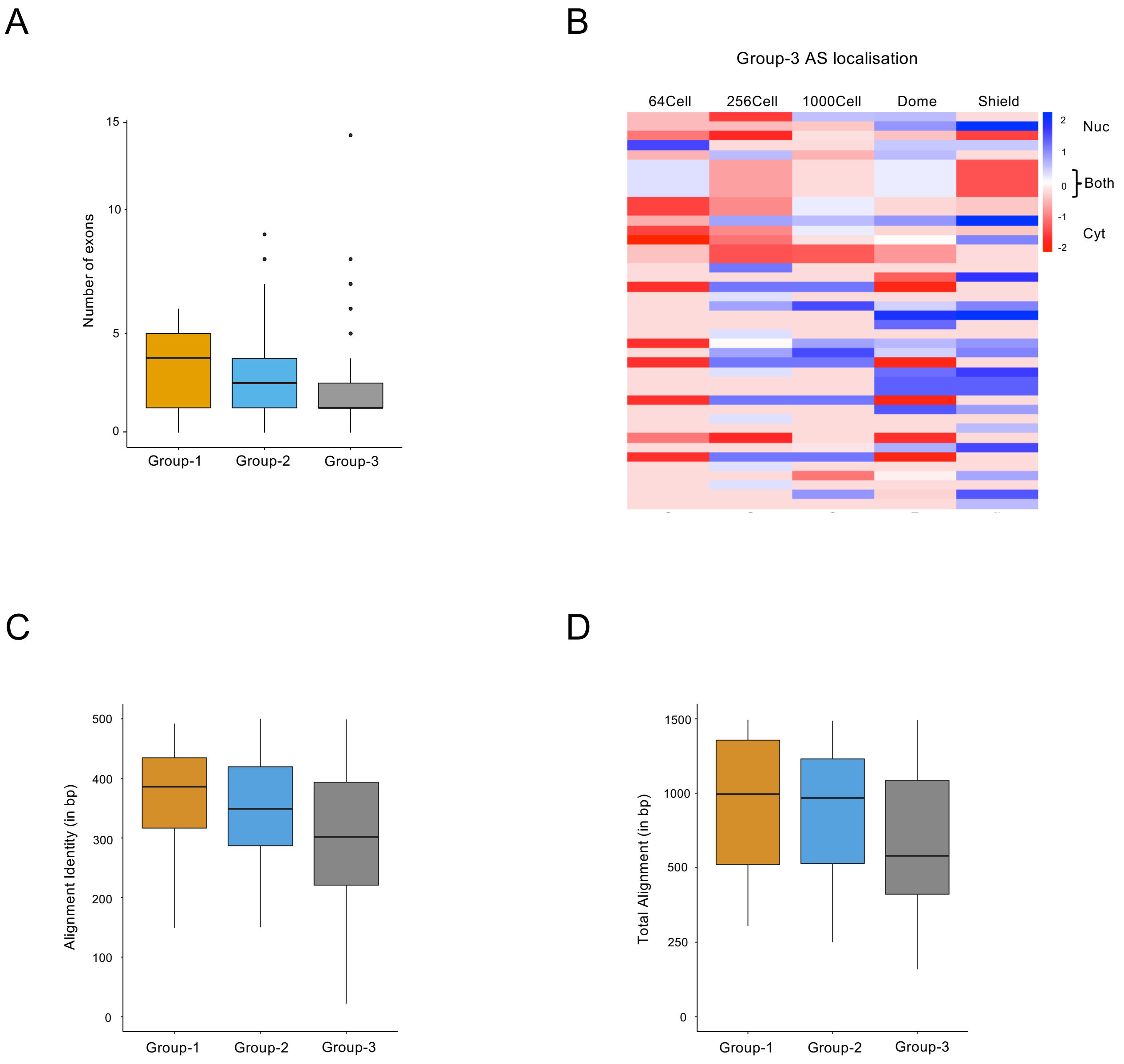


Supplementary Figure 5. (A) Boxplots of number of exons in the group-1, group-2 and group-3 AS. (B) Heatmap showing localization of group-3 AS in the nuclear (blue) and cytosolic fraction (red) with zebrafish development. The group-3 AS are equally distributed in the nuclear and cytosolic compartments. (C) Boxplots showing variation in alignment identity (bp) of group-1, group-2 and group-3 AS RNA with overlapping mRNA sequence. (D) Boxplots showing differences in total alignment (bp) of group-1, group-2 and group-3 AS RNA with their mRNA sequence. Both the graphs show higher alignment of group-1 AS with mRNA as compared to the other two groups.
